# FAN-C: A Feature-rich Framework for the Analysis and Visualisation of C data

**DOI:** 10.1101/2020.02.03.932517

**Authors:** Kai Kruse, Clemens B. Hug, Juan M. Vaquerizas

## Abstract

Chromosome conformation capture data, particularly from high-throughput approaches such as Hi-C and its derivatives, are typically very complex to analyse. Existing analysis tools are often single-purpose, or limited in compatibility to a small number of data formats, frequently making Hi-C analyses tedious and time-consuming. Here, we present FAN-C, an easy-to-use command-line tool and powerful Python API with a broad feature set covering matrix generation, analysis, and visualisation for C-like data (https://github.com/vaquerizaslab/fanc). Due to its comprehensiveness and compatibility with the most prevalent Hi-C storage formats, FAN-C can be used in combination with a large number of existing analysis tools, thus greatly simplifying Hi-C matrix analysis.

## INTRODUCTION

The development over the last decade of high-throughput techniques to study the three-dimensional organisation of the genome (Lieberman-Aiden et al. 2009; de Wit and de Laat 2012; Denker and De Laat 2016) in the nucleus has fuelled the characterisation of chromatin conformation in a wide variety of biological systems. These range from the organisation of the bacterial nucleoid (Le et al. 2013), to the *in vitro* characterisation of the molecular mechanisms that govern chromatin organisation in eukaryotes (Gassler et al. 2017; Haarhuis et al. 2017; Busslinger et al. 2017; Rao et al. 2017; Sanborn et al. 2015; Rhodes et al. 2020), reviewed in (Stadhouders et al. 2019), how this organisation is dynamically regulated during cell cycle (Naumova et al. 2013; Gibcus et al. 2018), development and differentiation (Hug et al. 2017; Flyamer et al. 2017; Du et al. 2017; Ke et al. 2017; Bonev et al. 2017; Chen et al. 2019), reviewed in (Hug and Vaquerizas 2018), and how it is affected in disease (Lupiáñez et al. 2015; Franke et al. 2016; Díaz et al. 2018), reviewed in (Spielmann et al. 2018).

Given the fundamental role that the correct organisation of chromatin in the nucleus plays for proper cell physiology, there is a growing need to integrate chromatin contact data in current studies examining different aspects of gene and genome regulation. Different techniques have been developed to study chromatin conformation at the single cell or population level, *in situ* Hi-C being the primary method of choice for analysing chromatin conformation in cell populations (Rao et al. 2014), reviewed in (Kempfer and Pombo 2019).

The large amounts of Hi-C data and increasingly specialised research questions have led to the development of diverse Hi-C analysis tools. Typically, these fall into one, rarely multiple of the following categories: Hi-C matrix generation, feature analysis, and visualisation (Pal et al. 2019; Ay and Noble 2015; Ing-Simmons and Vaquerizas 2019). Hi-C matrix generation tools convert FASTQ data from a Hi-C experiment into a normalised matrix of interaction strengths between pairs of genomic regions, accounting for false-positive interactions in the process. Feature analysis tools act on the Hi-C matrix to derive measures, models, and statistics that answer specific biological questions, such as the identification of topologically associating domains (Dixon et al. 2012; Sexton et al. 2012; Nora et al. 2012) and chromatin loops (Varoquaux et al. 2014; Rao et al. 2014), the 3D modelling of the chromatin fibre (Le Dily et al. 2017; Lin et al. 2019), or the identification of differential contacts between samples. Visualisation tools then enable the static display, and sometimes interactive exploration of the Hi-C matrix, and generally also of associated genomic data (Yardimci and Noble 2017; Ing-Simmons and Vaquerizas 2019).

The complexity of handling Hi-C data, owed in part to the vast amounts of data produced, prompted the development of several dedicated Hi-C storage formats in the form of compressed binary (Durand et al. 2016b) or hierarchical (Abdennur and Mirny 2019) files, or as text file specifications. The combination of specialised tools and available Hi-C formats results in a fragmentation of Hi-C analysis methods, which in turn causes a significant overhead for researchers analysing Hi-C data (Table 1).

**Table 1.**
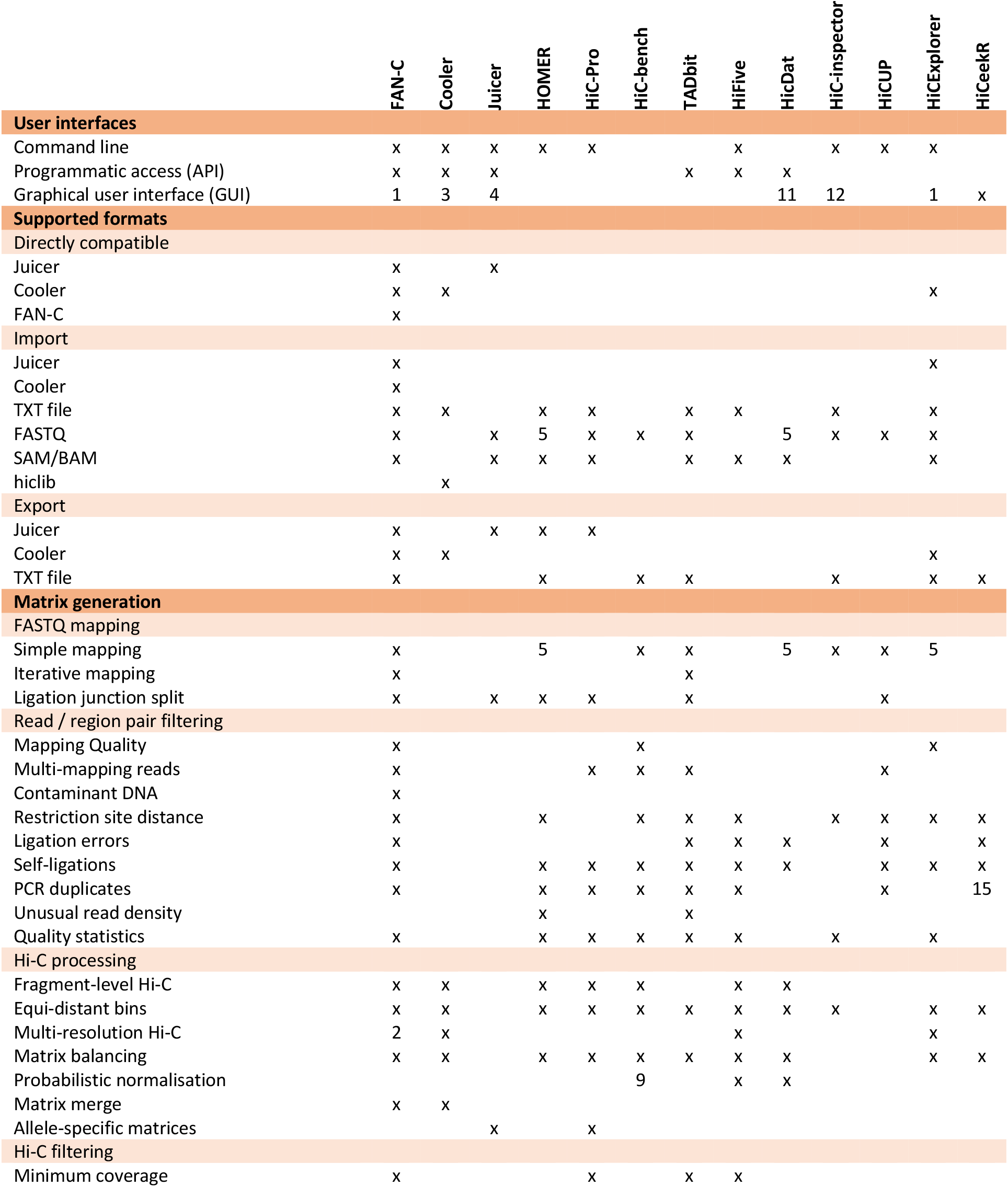
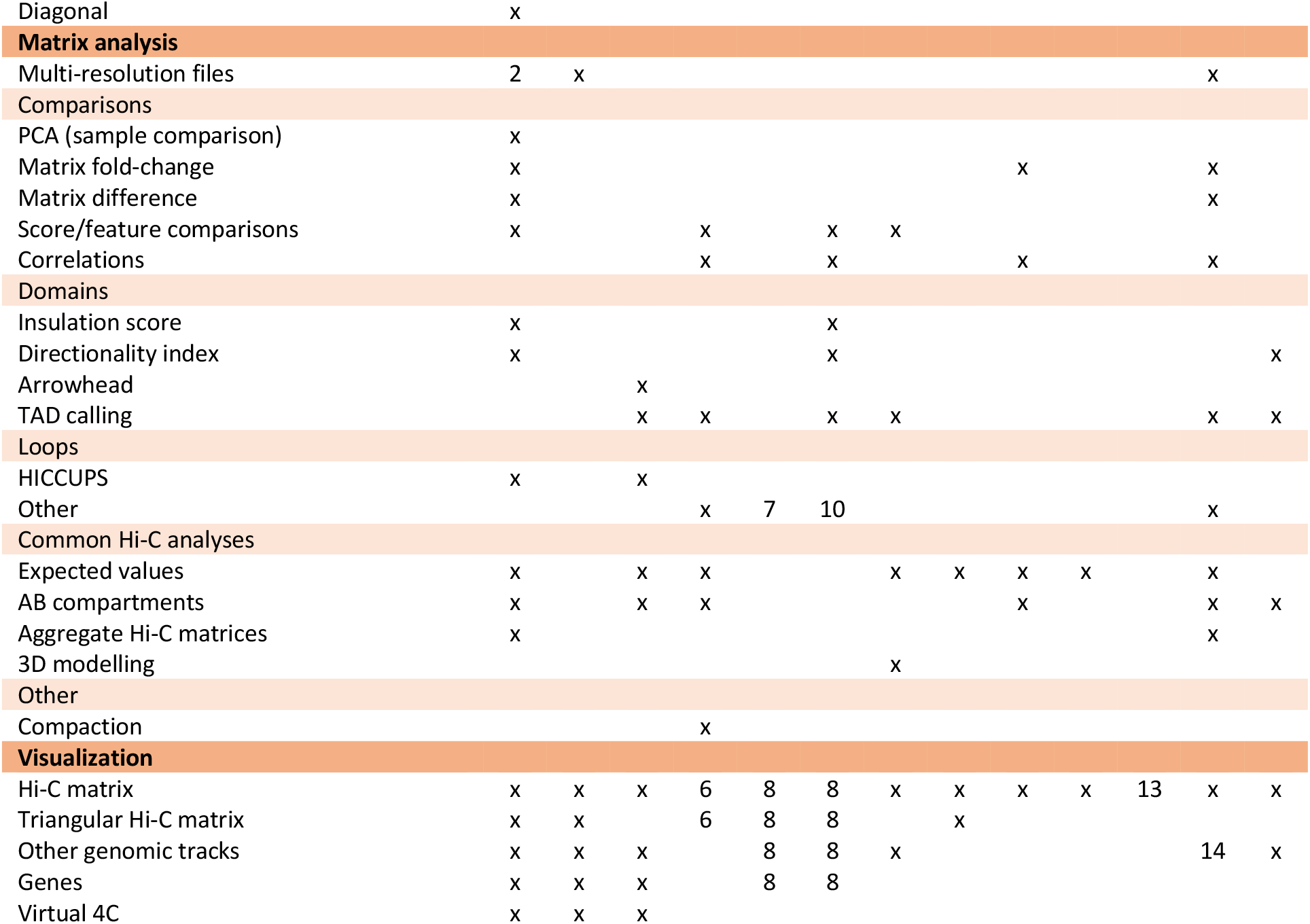
Feature comparison of different Hi-C analysis tools. Tools included in the comparison are Cooler (Abdennur and Mirny 2019) / HiGlass (Kerpedjiev et al. 2018), Juicer (Durand et al. 2016b) / Juicebox (Durand et al. 2016a), HOMER (Heinz et al. 2010), HiC-Pro (Servant et al. 2015), HiC-bench (Lazaris et al. 2017), TADbit (Serra et al. 2017), HiFive (Sauria et al. 2015), HicDat (Schmid et al. 2015), HiCInspector (Castellano et al. 2015), HiCUP, HiCExplorer (Ramírez et al. 2018), HiCeekR (Di Filippo et al. 2019). 1: Only for interactive plotting; 2: Support for Juicer multi-resolution files, but no native support; 3: In conjunction with HiGlass; 4: In conjunction with Juicebox; 5: Provides instructions for mapping, but no dedicated command; 6: Visualisation through Treeview; 7: With export for Fit-Hi-C; 8: Through compatibility with HiCPlotter; 9: Via HiCNorm; 10: Basic interaction enrichment; 11: Only pre-processing; 12: For interactive visualisation; 13: SAM/BAM visualisation through SeqMonk; 14: Limited support for other tracks, such as TAD separation score; 15: Only when previously marked in BAM file.

Here, we present FAN-C, a <u>F</u>ramework for the <u>AN</u>alysis of <u>C</u>-like data, an easy-to-use command-line tool and powerful Python API with a broad feature set covering matrix generation, analysis, and visualisation (Fig. 1). FAN-C uses a custom hierarchical storage format optimised for fast matrix access and common Hi-C matrix transformations. In addition, it is natively compatible and inter-convertible with the widespread Cooler (Abdennur and Mirny 2019) and Juicer (Durand et al. 2016b) Hi-C file formats, and can import a large variety of different text-based matrix inputs, such as those defined by HiC-Pro (Servant et al. 2015) and the 4D Nucleome project (Dekker et al. 2017). FAN-C includes a fully automated FASTQ-to-matrix pipeline, which can be adapted to the complexities and individual requirements of each specific Hi-C analysis, such as different species or analysis parameters. FAN-C also allows running each pipeline step individually, each with numerous customisation options. In addition, due to its broad file format support, FAN-C has the potential to integrate seamlessly with other tools, thereby significantly simplifying existing Hi-C analysis pipelines.

**Figure 1.**
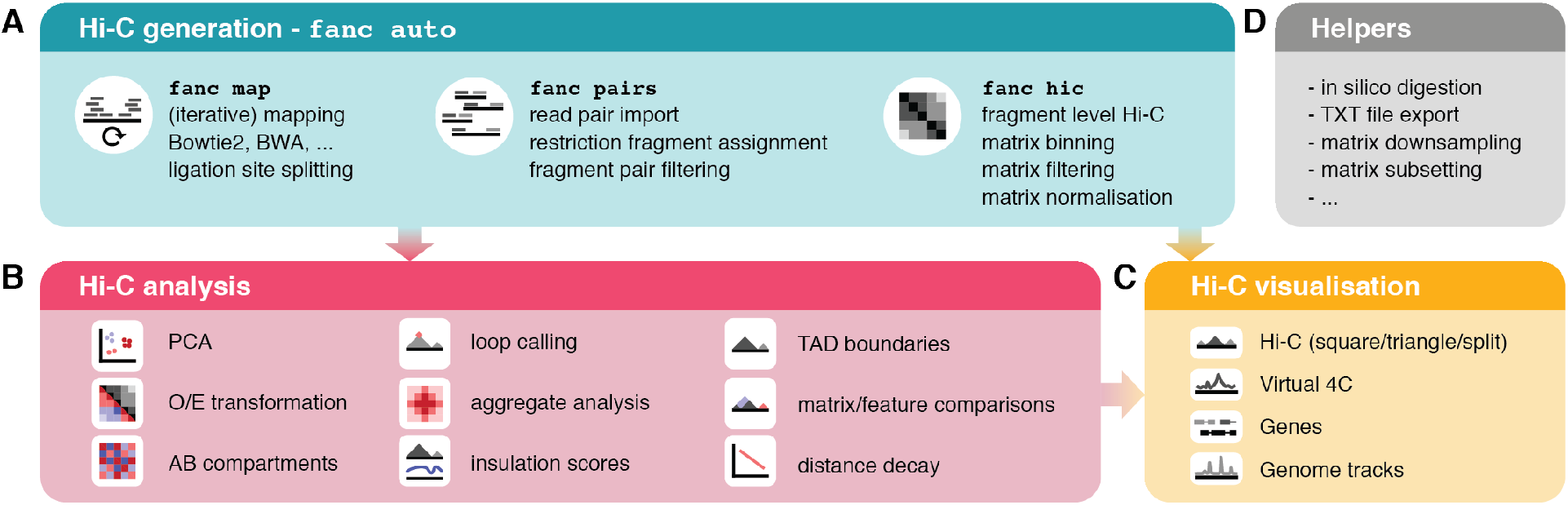
Overview of FAN-C functionality. A. Matrix generation features. B. Hi-C matrix analysis features. C. Hi-C visualization features. D. Helper tools

## RESULTS

### Hi-C matrix generation: from raw sequencing output to chromatin contacts

The first component of the FAN-C analysis framework consists of tools for matrix generation (Fig. 1A). This encompasses the mapping of sequencing reads to a reference genome, assignment of mapped reads to restriction fragments and the formation of interacting fragment-pairs, assembly of a fragment-level Hi-C matrix, and binning, as well as normalising that matrix at different resolutions. At each step, false-positive contacts need to be carefully filtered out in order to prevent matrix artefacts.

The primary tool for matrix generation in FAN-C is a fully automated pipeline, executable by a single command: `fanc auto`. It accepts a variety of automatically recognised input formats, including: i) unmapped reads in paired-end, optionally gzipped FASTQ files; ii) mapped reads from SAM or BAM files; and, iii) pre-processed read pairs or genomic contacts from other Hi-C pipelines in the form of text files (Fig. 2A-C). FASTQ files are mapped independently to a reference genome using either Bowtie2 or BWA-the choice of mapper is detected automatically from the genome index specified. To boost mapping efficiency, FAN-C can automatically detect and split reads at Hi-C ligation junctions, which are created by the cutting and religation of restriction sites. Further improvements to mapping efficiency can be achieved by enabling iterative mapping (Imakaev et al. 2012), where unaligned reads are truncated by a small number of base pairs and then attempted to align again (Fig. 2A).

**Figure 2.**
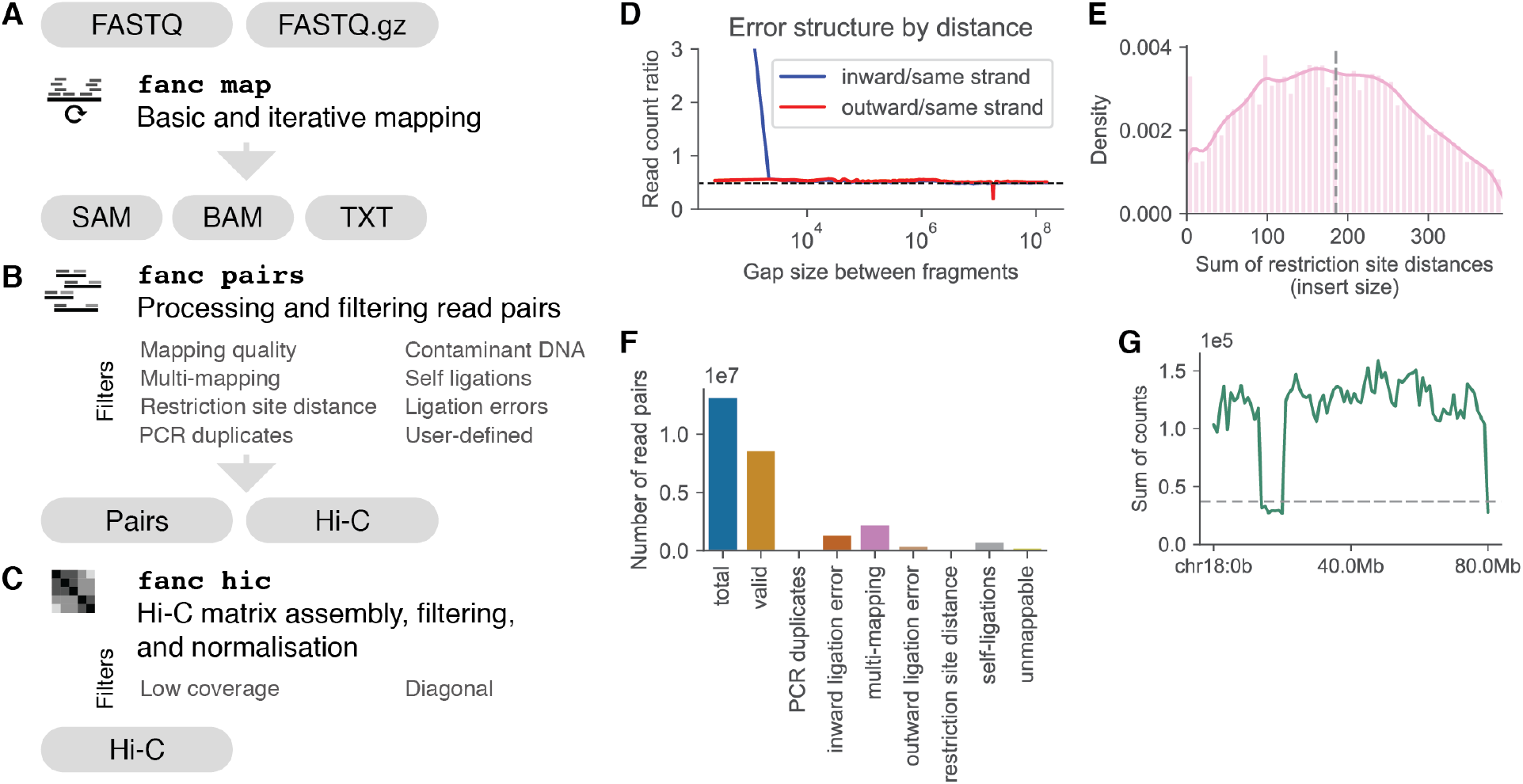
FAN-C matrix generation. A-C. Schematic overview of the matrix generation pipeline. A. Mapping features. B. Processing and filtering of Hi-C read pairs. C. Assembly, filtering and normalization of the Hi-C matrix from valid read pairs. D-F. FAN-C statistics plots using data from HUVEC Hi-C (Rao et al. 2014). D. Ligation error plot as in (Jin et al. 2013; Cournac et al. 2012). Dashed line indicates expected values. E. Density plot of the sum of restriction site distances (insert size) measured from the mapping location of a read to the nearest restriction site. Dashed line indicates median insert size. F. Summary statistics plot showing the read pairs removed by various filters. G. Coverage plot of a Hi-C matrix binned at 1kb resolution. Dashed line indicates the chosen coverage cutoff at 25% median coverage.

Mapped reads are then paired and assigned to restriction fragments (Fig. 2B). These are computed automatically using the restriction enzyme name and genome FASTA files, or can alternatively be supplied via a custom restriction map. Read pairs are filtered for common biases, including, among others, mapping quality, PCR duplicates, different types of ligation errors (Fig. 2D), and unexpected insert sizes (Fig. 2E). The filtering is highly customisable with a large selection of available filters, as well as the option to define custom filters using the Python API (Fig. 2B). Diagnostic plots with filtering statistics are generated automatically and are useful to inform the user about potential issues regarding the quality of Hi-C library or the set of parameters chosen for filtering (Fig. 2F).

Valid pairs, i.e., those that have passed the filtering steps above, are assembled into a fragment-level Hi-C matrix, which in turn is binned at various, customizable resolutions. Each binned matrix then undergoes a second round of filtering at the matrix level, including filters for low coverage of matrix bins (Fig. 2G), and is finally corrected for experimental and computational biases using Knight-Ruiz matrix balancing (Knight and Ruiz 2013) or, optionally, iterative correction (Imakaev et al. 2012) (Fig. 2C). Importantly, matrix rows and columns whose contact frequencies sum up to zero are explicitly ignored (or in some cases optionally imputed) by all FAN-C analysis methods. This avoids downstream analysis artefacts from falsely treating corresponding regions as if they had a complete lack of contacts, e.g. regions with poor mappability.

One of the key features of FAN-C is the ability to run each pipeline step independently, using dedicated commands. This enables the user to evaluate various parameter settings, and to perform parameter sweeps to test the robustness and ensure consistency of their analyses. Importantly, parameter changes can be made after the initial matrix generation, once bias statistics are available and a binned matrix can be investigated, without having to rerun the most time-consuming steps of Hi-C matrix assembly.

In order to maximise inter-compatibility with existing pre-processing, analysis, and visualisation pipelines, FAN-C includes several conversion tools. Valid pairs can be converted to Juicer’s Hi-C format using `fanc to-juicer`. Similarly, binned FAN-C matrices can be exported to multi-resolution Cooler files using `fanc to-cooler`, which are then compatible with cooltools (Venev et al. 2019) and HiGlass (Kerpedjiev et al. 2018) for visualisation.

### Matrix analysis: Chromatin compartments

FAN-C includes implementations of the most established analyses and measures for the characterisation of Hi-C matrix properties (Fig. 1B, Fig. 3A). Contact strength and the preference of contacts between certain genomic regions are particularly useful measures for gaining a global view of chromatin organisation. FAN-C implements several tools for this type of analysis:

i. Contact distance decay plots: the average contact strength between loci separated by a certain distance, also called “expected contacts”, is typically shown in a log-log plot of expected contacts *vs* distance (Fig. 3C). The slope and shape of the curve can inform about compaction of chromatin at various distance scales (Lieberman-Aiden et al. 2009);
ii. Observed/expected (O/E) transformation: a central transformation used by many analyses in which each pixel represents the (log2-)fold-change enrichment over the expected contact intensity for a region at that distance (Fig. 3D). Expected values are stored by FAN-C inside each matrix, allowing a fast, dynamic conversion of normalised into O/E contacts for various applications;
iii. Correlation matrices: the O/E matrix can further be transformed into a correlation matrix, in which each pixel i, j is calculated as the Pearson correlation coefficient between contacts in row i with column j (Fig. 3E, top). This highlights similarities and differences in contact profiles between loci, and reveals the partitioning of regions into the so-called A and B compartments in a plaid-like pattern (Lieberman-Aiden et al. 2009). Computationally, these are assigned using the sign of the correlation matrices’ first eigenvector (EV) (Fig. 3E, bottom). Due to the nature of EVs, positive entries do not necessarily correspond to the A, and negative to the B compartment. FAN-C offers the option to integrate information from a genomic FASTA file, which utilises the fact that the A compartment typically contains more GC-rich regions (Lieberman-Aiden et al. 2009) to flip the EV entry signs accordingly. The magnitude of the EV entry corresponding to a region is a rough measure for the region’s activity (Lieberman-Aiden et al. 2009; Flyamer et al. 2017);
iv. Saddle-plots: this helpful analysis allows the visualisation of interactions between A/B compartments of varying strength (Fig. 3F, top). To perform this analysis, regions are ordered and binned by their compartmentalisation strength (their entry in the correlation matrix EV) (Fig. 3F, bottom). The O/E values between regions of varying compartment strength provide a useful illustration of A and B segregation, and can further be used to quantify the level of compartmentalisation in the whole genome (Flyamer et al. 2017). A plot of cutoffs used for binning of regions is shown underneath the saddle plot (Fig. 3F, bottom). Unusually high or low EV entries, resulting, for example, from noisy or low mappability regions, can cause artefacts in the saddle plot, and are thus easily identifiable.

**Figure 3.**
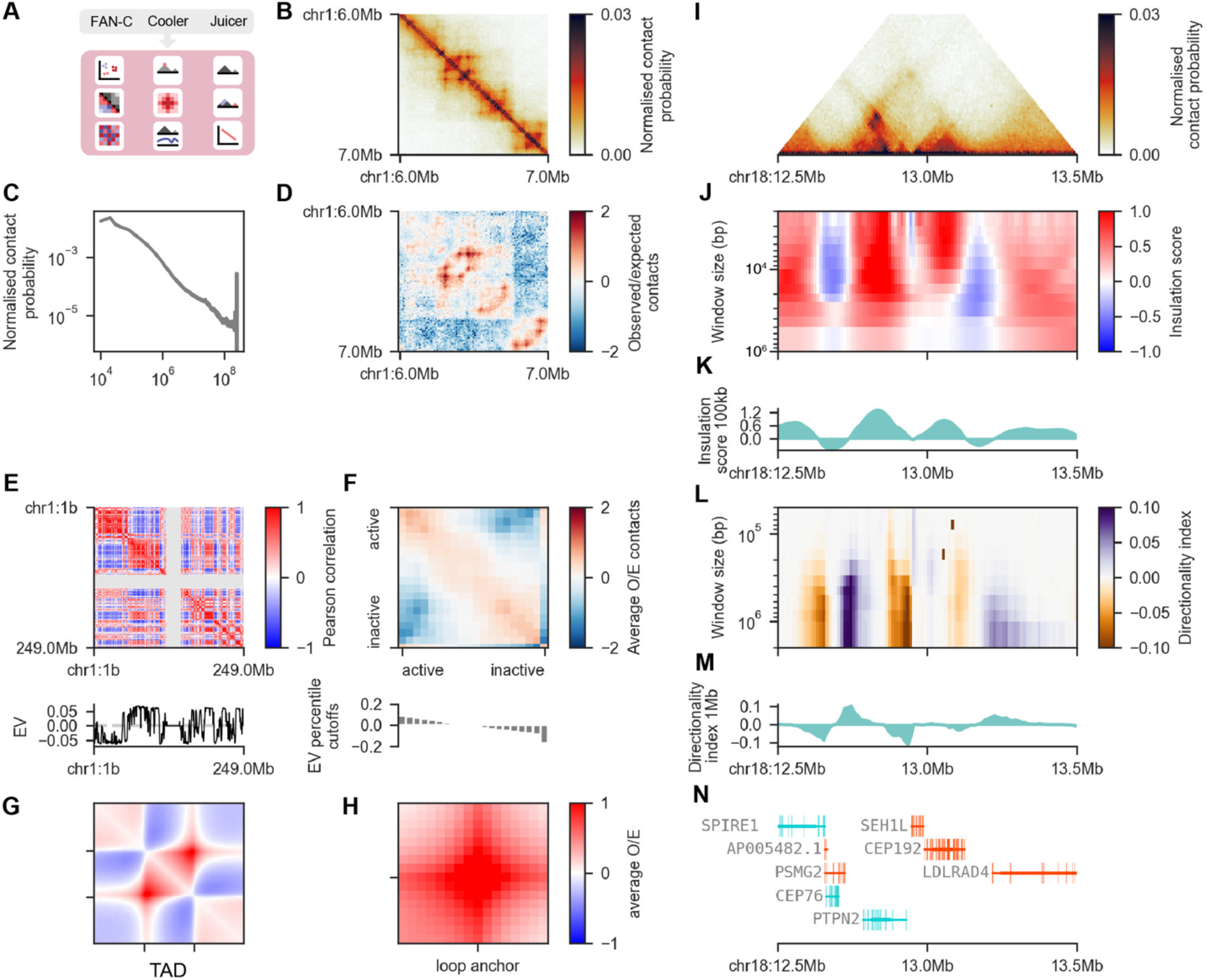
FAN-C analysis features. All analyses performed on GM12878 cells (Rao et al. 2014) on the 10kb resolution matrix, unless otherwise noted. A. Schematic representation of the analysis types available for FAN-C, Cooler, and Juicer matrices. B. Hi-C matrix plot of a sample region with 10kb resolution. C. Log-log “Distance decay” plot of the expected normalised contact frequency against locus distance. D. Log2-observed/expected (O/E) matrix for the same region as in A. E. 500kb resolution correlation matrix / A/B compartment plot of chromosome 1 (top) and its first eigenvector (EV) (bottom). F. “Saddle plot” showing preferential interactions of active/active and inactive/inactive regions (top), and bar plot showing the cutoffs used for binning regions by the corresponding EV entry magnitude (bottom). Note the outlier on the far right. G. Aggregate TAD plot showing the average log2-O/E in and around arrowhead domains (Rao et al. 2014). H. Aggregate loop plot showing the average log2-O/E at peaks called by HICCUPS (Rao et al. 2014). I-N. Example region on chromosome 18 highlighting additional analyses available in FAN-C and the possibility of “genome browser” style plotting. I. Triangular Hi-C matrix plot. J. Heatmap showing insulation scores calculated using different window sizes. K. Insulation score track for a window size of 100kb. L. Heatmap showing directionality index results for multiple window sizes. M. Directionality index track for a window size of 1Mb. N. Gene plot using data from Gencode (v19) (Harrow et al. 2012).

### Matrix analysis: TADs, chromatin loops, and aggregate analysis

High-resolution analyses of Hi-C matrices have revealed conserved matrix features that appear to be common across higher eukaryotes. These include topologically-associating domains (TADs) (Dixon et al. 2012; Nora et al. 2012; Sexton et al. 2012), regions of increased self-interaction that are separated by insulating boundaries from neighbouring domains and are visible as squares in a Hi-C matrix (Fig. 3B), and chromatin loops (Rao et al. 2014), enriched discrete contacts between pairs of regions that show up as local areas of increased contact intensity in the matrix (Fig. 3B). FAN-C contains implementations of the most widely used algorithms for TAD and loop analysis:

i. Insulation score and directionality index: Genomic regions between TADs, characterised by their strong insulating effect on neighbouring domains, can be identified using the insulation score (Crane et al. 2015) (Fig. 3J and K), or the directionality index (Dixon et al. 2012) (Kruse et al., 2016) (Fig. 3L and M). The resulting insulation tracks, quantifying the insulating effect of each region, can be exported to a range of established genomic formats, so they can easily be imported into genome browsers or used in other analysis pipelines.
ii. Chromatin loops: discrete peaks in the Hi-C matrix correspond to loops between genomic regions (Rao et al. 2014). To identify these loops, FAN-C includes a CPU implementation of HIC-CUPS, a local-neighbourhood based loop calling algorithm (Rao et al. 2014), which can be parallelised on a computational cluster.
iii. Aggregate plots: to help with the identification of global trends across chromatin contact datasets, a genome-wide overview of the conformation around TADs, loops, or other genomic features such as promoters, can be obtained with aggregate plots, which represent an average conformation around all regions of interest (Flyamer et al. 2017). FAN-C implements the generation of aggregate plots from any list of regions or region pairs, with useful presets for TAD (Fig. 3G) and loop (Fig. 3H) aggregate plots. The aggregation process and the look of the aggregate matrix plot are highly customisable, for example by controlling size and resolution of the matrix, as well as colours and annotations of the final plot.

### Matrix comparison: highlighting and identifying differential features

A central task in Hi-C matrix analysis is the comparison of multiple datasets. A number of tools have been developed to identify and quantify differences between Hi-C matrices (Heinz et al. 2010; Stansfield et al. 2018; Lun and Smyth 2015; Ardakany et al. 2019; Djekidel et al. 2018). FAN-C focusses on the representation and visualisation of differences, and can therefore function as a direct extension to existing approaches.

Principal component analysis (PCA) can provide a general overview of the similarity of several datasets by performing a pairwise comparison of matrix entries (Hug et al. 2017; Díaz et al. 2018). FAN-C implements methods for PCA analysis of Hi-C contacts, such as for Hi-C library replicates and samples. Since not all pixels in a matrix are equally informative, e.g., regions far away from the diagonal or inter-chromosomal contacts are often dominated by noise, FAN-C includes a number of filters, such as distance between loci, or largest variance between samples, to only consider the most informative contacts in a matrix.

Side-by-side comparison of matrices or measures derived from these are widely used and can be very useful in displaying changes in chromatin contacts in a visual manner (Fig. 4A). However, features specific to only one matrix can often be more effectively highlighted by calculating matrix differences (Fig. 4B) or fold-changes (Fig. 4C). FAN-C implements functions to calculate differences and fold-changes on matrices and associated tracks, such as the insulation score (Fig. 4D and E) or compartment strength. Users can also supply external tracks in a compatible genomic format (BED, GFF, BigWig) for comparison. The resulting tracks and matrices can be used as input to any FAN-C function in the same fashion as regular objects, including the following visualisations.

**Figure 4.**
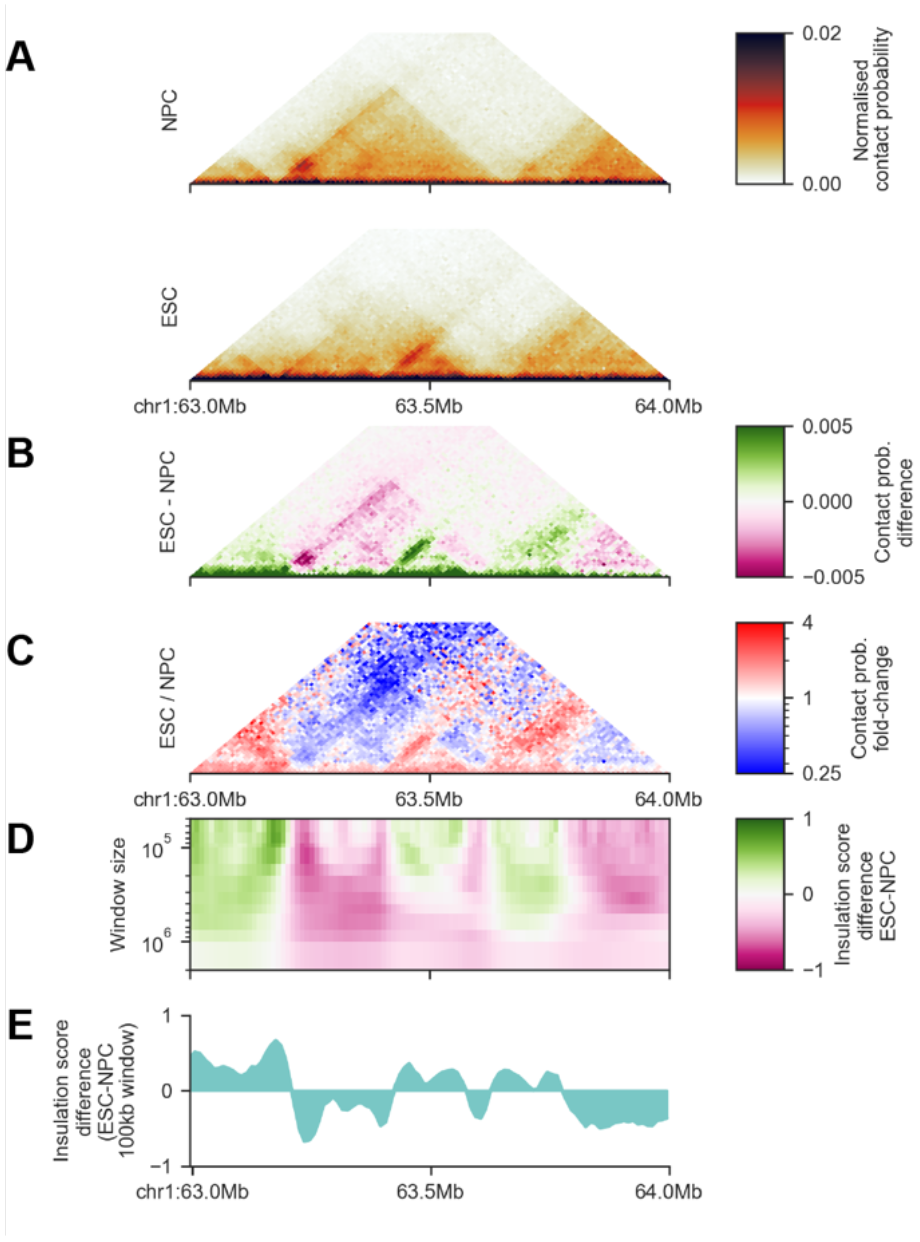
FAN-C Matrix and feature comparisons. A-E. Comparison of mouse embryonic stem cell (ESC) and neural precursor cell (NPC) Hi-C matrices at 10kb resolution (Bonev et al. 2017). A. Triangular Hi-C matrix plot of ESC and NPC of a sample region on chromosome 1. B. Difference matrix obtained by pixel-wise subtraction of contacts in NPC from contacts in ESC. C. Fold-change matrix obtained by pixel-wise division of contacts in NPC by contacts in ESC. D. Heatmap of insulation score differences between ESC and NPC at various window sizes. E. Insulation score difference track using a 100kb window.

### Plotting: interactive and publication-ready visualisation of Hi-C and related data

FAN-C includes an implementation of an advanced yet easy to use plotting library for C-derived datasets (Fig. 1C). A number of diagnostic plots are generated as part of the `fanc auto` command, including filtering statistics for read pairs, biases in ligation frequency, and chromosomal coverage. Specific versions of the plots can also be produced individually, to allow for a thorough comparison of parameters used in an analysis. Plots related to Hi-C matrix-derived measures, such as correlation matrix, saddle, and aggregate plots (Fig. 3E-H) are part of the individual analysis functions. Plots for time-consuming analyses, such as aggregating matrices over a large number of regions, can easily be tweaked and adjusted without having to re-compute the entire analysis.

In addition to static plots, FAN-C also includes a basic interactive genome browser that allows for the interactive browsing of Hi-C and additional genomic datasets. These include various different representations of Hi-C matrices: square (Fig. 3B); triangular (Fig. 3I); mirrored, in which two triangular Hi-C matrices are shown above and below a horizontal dividing line; and “split”, where the diagonal separates two different matrices in a square plot. A slice of a Hi-C matrix can also be visualised as a virtual 4C plot, which shows the strength of contacts between a specific genomic region and a genomic interval, as a line plot. This can be useful, for example, to visualise specific pairwise interactions, or even to detect genomic rearrangements such as translocations (Díaz et al. 2018) or genome insertions (Kruse et al. 2019). All of the above matrix plots can also be used to display difference (Fig. 4B) and fold-change (Fig. 4C) maps.

Several plot types are available for region-based data in a standard genomic data format, including support for BED, GFF, BigWig, and Tabix-indexed files. These can be displayed as boxes coloured by strand, optionally grouped into layers by a user-defined attribute, or - in case they contain scores - as bar or line plots. Insulation score and directionality index results, which depend on a chosen window size parameter, have a dedicated plot type that visualises scores for multiple window sizes simultaneously in a heatmap (Fig. 3J and K, Figure 4D), similar to the previously suggested “domainogram” (De Wit et al. 2008). Finally, genome annotations can be plotted with intron/exon visualisations, as well as depicting strand information (Fig. 3N).

In addition to interactive visualisation, FAN-C includes a powerful plotting API for generating vector-based, publication-ready visualisations. Each type of interactive plot outlined above is also available through the API, and is individually customisable. Since it is based on the major Python plotting library matplotlib, it is easily extensible and can easily be integrated in existing plotting scripts. As a demonstration, everything in Fig. 3 and 4 of this manuscript, apart from annotations and schematics, has been generated entirely using the FAN-C plotting API. This makes FAN-C not only useful for Hi-C matrix analysis, but also for users wanting to produce high-quality plots from pre-computed matrices to integrate alongside their existing visualisations.

## CONCLUSIONS

Here we introduce FAN-C as an open-source, versatile, flexible, and powerful tool for Hi-C analysis. FAN-C is bundled with an extensive documentation, available at https://vaquer-izaslab.github.io/fanc, and sample datasets at https://github.com/vaquerizaslab/fanc. The documentation includes detailed examples of how to use the command line tools and, for advanced applications, the versatile Python API. When designing FAN-C functionality, we have specifically tried to include the most widely-used measures and analyses with sensible defaults, while offering fine-grained control over analysis details. A side-by-side comparison with existing Hi-C analysis tools shows the broad spectrum of analysis options covered by FAN-C (Table 1). Due to its feature set and compatibility with the most established Hi-C formats, we envisage FAN-C to occupy a central position in many Hi-C pipelines.

## AVAILABILITY

FAN-C is available at the Vaquerizas Laboratory GitHub page: https://github.com/vaquerizaslab/fanc

## AUTHOR CONTRIBUTIONS

Conceptualization, K.K, C.B.H. and J.M.V.; Methodology, K.K. and C.B.H.; Formal Analysis, K.K. and C.B.H.; Writing – Original Draft, K.K. and J.M.V.; Writing – Review and Editing, K.K., C.B.H. and J.M.V.; Funding Acquisition, J.M.V.; Supervision, J.M.V.

## ACKNOWLEDGEMENTS

We thank Noelia Díaz, Benjamín Hernández-Rodríguez, Elizabeth Ing-Simmons, and Jahnavi Bhaskaran from the Vaquerizas laboratory for helpful discussions, suggestions for implementation and beta testing of FAN-C. We are grateful to Alexis G. Grimaldi for help with early implementations of code. We thank Tom Sexton (IG-BMC, Strasbourg) for beta testing of the software. We are grateful to Elizabeth Ing-Simmons and Tom Sexton for providing comments on the manuscript.

